# A case for a reverse-frame coding sequence in a group of positive-sense RNA viruses

**DOI:** 10.1101/664342

**Authors:** Adam M. Dinan, Nina I. Lukhovitskaya, Ingrida Olendraite, Andrew E. Firth

**Affiliations:** Division of Virology, Department of Pathology, University of Cambridge, Tennis Court Road, Cambridge, CB2 1QP, UK

**Author notes:** **Correspondence to:** Andrew E. Firth.

## Abstract

Positive-sense single-stranded RNA viruses form the largest and most diverse group of eukaryote-infecting viruses. Their genomes comprise one or more segments of coding-sense RNA that function directly as messenger RNAs upon release into the cytoplasm of infected cells. Positive-sense RNA viruses are generally accepted to encode proteins solely on the positive strand. However, we previously identified a surprisingly long (~1000 codons) open reading frame (ORF) on the negative strand of some members of the family *Narnaviridae* which, together with RNA bacteriophages of the family *Leviviridae*, form a sister group to all other positive-sense RNA viruses. Here, we completed the genomes of three mosquito-associated narnaviruses, all of which have the long reverse-frame ORF. We systematically identified narnaviral sequences in public data sets from a wide range of sources, including arthropod, fungi and plant transcriptomic datasets. Long reverse-frame ORFs are widespread in one clade of narnaviruses, where they frequently occupy >95% of the genome. The reverse-frame ORFs correspond to a specific avoidance of CUA, UUA and UCA codons (i.e. stop codon reverse complements) in the forward-frame RNA-dependent RNA polymerase ORF. However, absence of these codons cannot be explained by other factors such as inability to decode these codons or GC3 bias. Together with other analyses, we provide the strongest evidence yet of coding capacity on the negative strand of a positive-sense RNA virus. As these ORFs comprise some of the longest known overlapping genes, their study may be of broad relevance to understanding overlapping gene evolution and *de novo* origin of genes.

## INTRODUCTION

Traditionally, viruses have been divided between seven Baltimore classes based on the nature of the nucleic acid of their genomes and their replicative intermediates. The seven classes are positive-sense, negative-sense and double-stranded RNA viruses, single-stranded and double-stranded DNA viruses, retroviruses and pararetroviruses (Baltimore 1971). Of these, the single-stranded positive-sense or (+)ssRNA viruses comprise the largest and most diverse group of eukaryote-infecting viruses (Dolja and Koonin 2011). The group includes many important human and animal pathogens (such as dengue, Zika, yellow fever, hepatitis C, foot-and-mouth disease, polio, chikungunya, SARS and MERS viruses) besides the majority of plant viruses.

To fully understand the molecular biology of viruses, it is crucially important to know their coding capacity. In recent years, a number of “hidden” coding open-reading frames (ORFs) have been discovered in the genomes of various (+)ssRNA viruses (Chung et al. 2008; Loughran et al. 2011; Fang et al. 2012; Firth 2014; Smirnova et al. 2015; Napthine et al. 2017; Lulla et al. 2019). Such “hidden” genes tend to be very short and/or to overlap previously known coding ORFs, explaining why they have escaped previous detection prior to the application of sensitive comparative genomic methods. However, all of these experimentally verified novel coding ORFs have been in the positive-sense. To our knowledge, no negative-sense coding ORF has *ever* been demonstrated in *any* (+)ssRNA virus. Thus, an unanswered question in virology is whether the negative strand of (+)ssRNA viruses can encode proteins.

In 2013, by acquiring and analyzing RNA transcriptomic datasets for several mosquito and other dipteran species, we identified two “mosquito-associated” narna-like viruses (Cook et al. 2013). Similar to other narnaviruses, one strand contains a single long ORF that encodes a protein inferred by homology to be the RNA-dependent RNA polymerase (RdRp). However, we were surprised to notice that both sequences also contain a reverse-frame ORF (rORF) covering nearly the entire sequence of ~3000 nt. We also identified related rORF-containing sequences in public *Puccinia striiformis* and *Uromyces appendiculatus* transcriptome shotgun assembly (TSA) data sets. The extreme divergences between these sequences and the mosquito-associated sequences (~22% amino acid identity in the RdRp sequence) effectively rule out a region of such length being preserved free of stop codons by chance, and thus we hypothesized that the rORF represented a *bona fide* protein-coding sequence.

As currently defined by the International Committee on Taxonomy of Viruses (ICTV), the family *Narnaviridae* contains two genera: *Mitovirus* and *Narnavirus*. Both genera contain single-stranded positive-sense RNA viruses which are non-encapsidated and hence are expected to be transmitted either vertically though cell division or horizontally during host mating. Mitoviruses replicate in the mitochondria of host cells whereas narnaviruses replicate in the cytoplasm. These viruses were originally described as infecting fungi, but related viruses have since been observed in transcriptomic data sets derived from diverse organisms. The narnaviral positive strand normally contains a single long ORF that covers most of the genome and encodes an RNA-dependent RNA polymerase (RdRp) which catalyses viral replication. The narnaviral RdRp is highly divergent from those of other eukaryotic RNA viruses, and shows closer homology to the RdRps of RNA bacteriophages in the family *Leviviridae* (Rodriguez-Cousiño et al. 1991; Esteban et al. 1992; Wolf et al. 2018). Detailed comparative genomic and phylogenetic analyses suggest that the *Narnaviridae* are descended from a levivirus-like bacteriophage which may have been carried within the bacterial progenitor of mitochondria at the point of eukaryogenesis, followed by loss of the capsid protein giving rise to capsidless (“naked”) RNA elements (Koonin et al. 2015; Wolf et al. 2018). The escape of a group of these viruses into the cytosol may have given rise to the *Narnavirus* genus (Koonin and Dolja 2014; Wolf et al. 2018).

The prototypical and best studied narnaviruses are the *Saccharomyces cerevisiae* 20S and 23S RNA viruses (ScNV-20S and ScNV-23S, respectively) (reviewed in Wickner et al. 2013). ScNV-20S persistently infects most laboratory strains of the yeast *Saccharomyces cerevisiae*, whereas fewer strains carry ScNV-23S. Their genomes have no 3′ poly(A) tail and it is not known if they have a 5′ cap structure (Rodríguez-Cousiño et al. 1998). The genomes do not encode capsids, but form ribonucleoprotein complexes with the RdRp in a 1:1 stoichiometry in the host cell cytoplasm; the RdRp interacts with both the 5′ and 3′ ends which may help protect the viral RNA from degradation by host exonucleases (Solórzano et al. 2000; Fujimura and Esteban 2004; Fujimura and Esteban 2007). Under suitably inducing conditions such as heat shock and nitrogen starvation, copy numbers of either virus can reach 100,000 copies per cell (Kadowaki and Halvorson 1971; Wejksnora and Haber 1978; Esteban et al. 1992). Approximately 98–99% of viral RNA in the cell is in a single-stranded positive-sense form, whereas the remainder exists as single-stranded negative-sense replication intermediates (Rodriguez-Cousiño et al. 1991; Fujimura et al. 2005). When cells are grown at high temperature, double-stranded forms (known as W for ScNV-20S and T for ScNV-23S) accumulate, but these appear to represent by-products and not replication intermediates (Wesolowski and Wickner 1984; Rodríguez-Cousiño et al. 1998; Fujimura et al. 2005).

To further investigate the presence of the rORF, and now that many more sequences are available in public sequence databases, in this work we present a comprehensive comparative analysis of narnaviral genomes, in which we assess the prevalence, distribution and sequence features of rORFs. Two major clades of narnavirus are identified, for which we propose the establishment of the genera *Alphanarnavirus* and *Betanarnavirus*, with the former clade containing all sequences with long rORFs. Overall codon usage is similar in alphanarnaviruses with and without long rORFs, but the former display a highly specific avoidance of CUA, UUA and UCA codons over large regions of the RdRp ORF, corresponding to an absence of stop codons in the rORF. We explore possible reasons for the avoidance of CUA, UUA and UCA and conclude that the most plausible explanation is selection to maintain an rORF, indicating that the rORF is functional.

## RESULTS

### Completion of narnaviral genomic sequences

We completed three genomic sequences for previously described rORF-containing narnaviruses (Cook et al. 2013), and used these genomes as a reference set for comparative analyses. The sequences with GenBank accession numbers KF298275.1, KF298276.1 and KF298284.1 were extended at their 5′ termini by 479 nt, 10 nt and 364 nt, respectively, and at their 3′ termini by 43 nt, 9 nt and 9 nt, respectively (**Figure 1A; Supplementary File S1**). The latter two sequences are the same length (3093 nt) and encode RdRps which are 97.95% identical at the amino acid level; hence, they were considered to represent a single viral species. Given that the samples from which these sequences were originally obtained were isolated from mosquitoes in the *Ochlerotatus* genus, we name the viruses Ochlerotatus-associated narna-like virus 1 (ONLV1; sequence KF298275.1) and ONLV2 (sequences KF298276.1 and KF298284.1) (**Figure 1A**). The AUG codons of the three RdRp ORFs each start at the 7th nucleotide of the respective sequences (contexts: GUC**AUG**A, GUU**AUG**G and GUU**AUG**G), the same position as that of the RdRp AUG in ScNV-23S (GenBank: NC_004050.1). The RdRp-encoding ORFs are 3045, 3075 and 3075 nucleotides in length, respectively. The RdRp stop codons (UGA, UAA and UAA) end at positions 14 nt, 13 nt and 13 nt, respectively, from the 3′ termini of the genomes. Hence, the length of the 3′ UTR mirrors closely that of ScNV-20S (12 nt; GenBank: NC_004051.1) but is shorter than that of ScNV-23S (59 nt).

**Figure 1.**
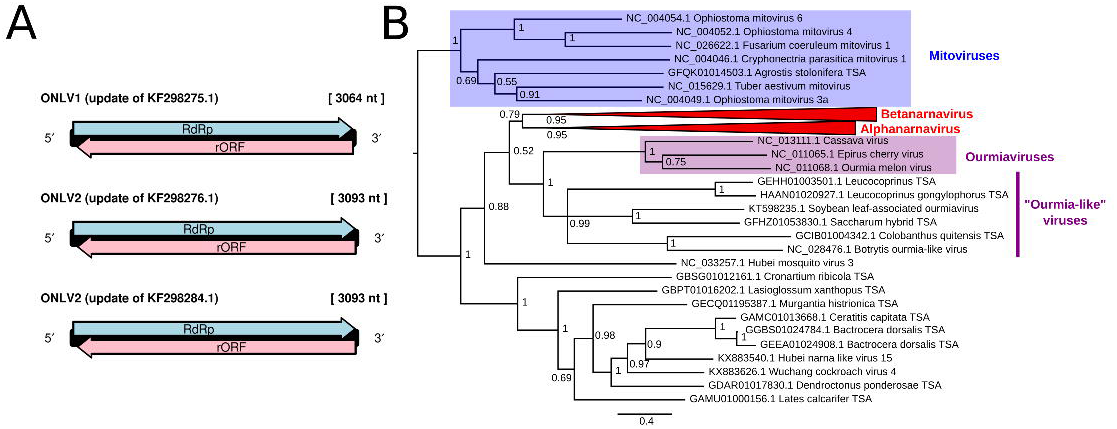
Narnavirus genome structure and taxonomy. (**A**) Updated narnaviral genome sequences: genome structures of ONLV1 and ONLV2. (**B**) Bayesian phylogeny of narnaviruses and selected outgroup taxa. Alphanarnaviruses and betanarnaviruses form clades with posterior probabilities of 0.95. The tree is rooted with the mitoviral clade as an outgroup. Accession numbers are for the nucleotide sequences from which the corresponding protein sequences were derived.

### An expanded narnaviral phylogeny

We employed tblastn (Camacho et al. 2009) and hmmsearch (Finn et al. 2011), using hidden Markov model based profiles, pHMM) to identify sequences encoding proteins closely related to narnaviral RdRps in the NCBI non-redundant (nr) and TSA databases. In total, 124 unique such sequences were identified – 46 in the nr database and 78 in the TSA database. The holobiont sources of these TSA sequences were phylogenetically diverse and included 21 arthropod, 8 fungal and 10 plant species. To place these sequences in context, we also selected a number of mitovirus and ourmiavirus sequences as outgroups, besides additional narna-like virus sequences from Shi et al. (2016), giving 141 sequences in total (**Supplementary Data Set S1**). We oriented all sequences (by reverse complementing if necessary) so that the RdRp ORF was on the forward strand.

To assess the relationships among sequences, an alignment of the predicted RdRp proteins was generated using MUSCLE (Edgar 2004) (**Supplementary File S2**), and a Bayesian Markov chain Monte Carlo (MCMC)-based tree was constructed from this alignment using MrBayes (Huelsenbeck and Ronquist 2001; **Figure 1B**). As expected, the free-floating genus *Ourmiavirus*, which includes plant-associated viruses with tripartite genomes, grouped unambiguously with the narnaviruses (Turina et al. 2017; Rastgou et al. 2009; Wolf et al. 2018), whereas mitoviruses formed a sister clade (**Figure 1B**). A relatively small, but highly diverse set of fungal- and arthropod-derived sequences clustered with the ourmiaviruses (**Figure 1B**). Recent evidence shows that these “ourmia-like” viruses display a range of genomic architectures, which can be segmented (as in the case of the *bona fide* ourmiaviruses) or non-segmented, occurring in mono- and di-cistronic forms (reviewed in Dolja and Koonin 2018). For example, botrytis ourmia-like virus, which appears to have a non-segmented genome, clusters unambiguously with ourmiaviruses rather than with the narnaviruses (Donaire et al. 2016; **Figure 1B**).

### Two major clades of narnaviruses

The narnaviral sequences predominantly fell into one of two major clades, both of which had Bayesian posterior probabilities of 0.95 (**Figure 1B**). Based on these data, we propose that the genus *Narnavirus* be subdivided to form two new genera, which we name *Alphanarnavirus* and *Betanarnavirus*, besides additional unclassified sequences.

The alphanarnaviral clade contains the prototypical narnaviruses, ScNV-20S and ScNV-23S, as well as sequences associated with a range of other fungal taxa, including members of the divisions Ascomycota, Basidiomycota and Entomophthoromycota, and the divergent subphylum Mucoromycotina (**Figure 2**). The alphanarnaviral sequences ranged in length from 1805 nt to 3874 nt (**Supplementary Figure S1**), and the pairwise amino acid identities of the corresponding set of RdRps ranged from 16.8% to 99.8% (**Supplementary File S3)**. Putative viral sequences containing an rORF all clustered within the alphanarnaviral clade, and none in the betanarnaviral clade. However, the rORF-containing sequences appear not to form a monophyletic clade, but instead cluster in several regions of the phylogeny, and are found in sequences derived from fungi, arthropods and plants (**Figure 2: red bars**). The core RdRp catalytic regions – motifs A to E in the palm domain and motifs F and G in the fingers (Wu et al. 2015; te Velthuis 2014) – are well-conserved despite the overall high degree of sequence divergence (**Supplementary Figure S2**).

**Figure 2.**
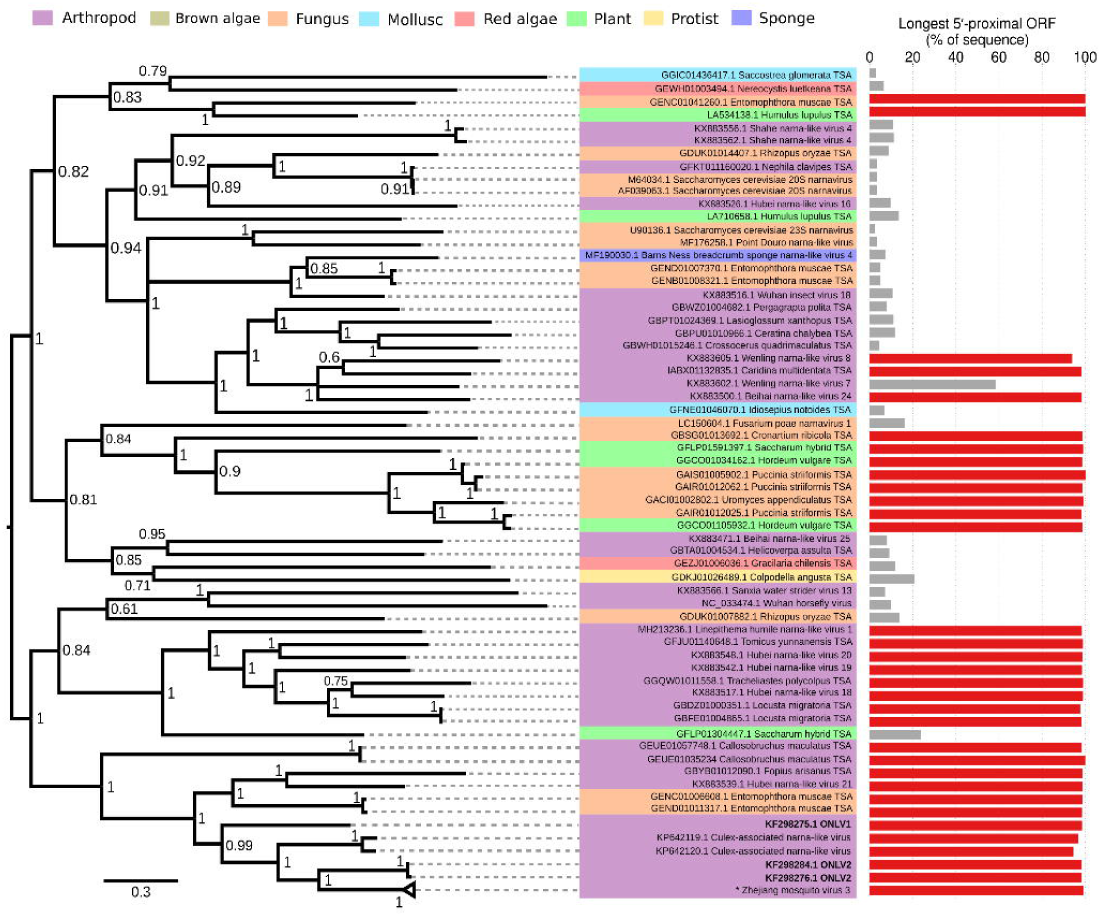
Midpoint-rooted Bayesian phylogenetic tree of alphanarnaviruses. The longest 5′-proximal ORF in the negative-strand R0 frame is shown in the bars to the right. Taxa with R0-frame ORFs occupying over 90% of the sequence are indicated with red bars. Nine highly similar sequences for Zhejiang mosquito virus 3 (indicated with an asterisk) are collapsed to a single taxon. Accession numbers are for the nucleotide sequences from which the corresponding protein sequences were derived.

The betanarnaviral clade includes several pathogens of unicellular eukaryotes, including the oomycete-infecting Phytophthora infestans RNA virus 4 (PiRV4; Cai et al. 2012) and the protozoan-associated Leptomonas seymouri narna-like virus 1 (Lye et al. 2016), as well as sequences from red algae, brown algae, myxozoa and arthropods (**Figure 3**). The betanarnaviral sequences ranged in length from 2215 nt to 3610 nt (**Supplementary Figure S1**), and the pairwise RdRp amino acid sequence identities ranged from 15.2% to 99.9% (**Supplementary File S4)**. In the case of Leptomonas seymouri narna-like virus 1, the genome may be bipartite, with the RdRp being encoded on the longer (L) segment, although the functional association of the putative segments has not been shown experimentally (Lye et al. 2016).

**Figure 3.**
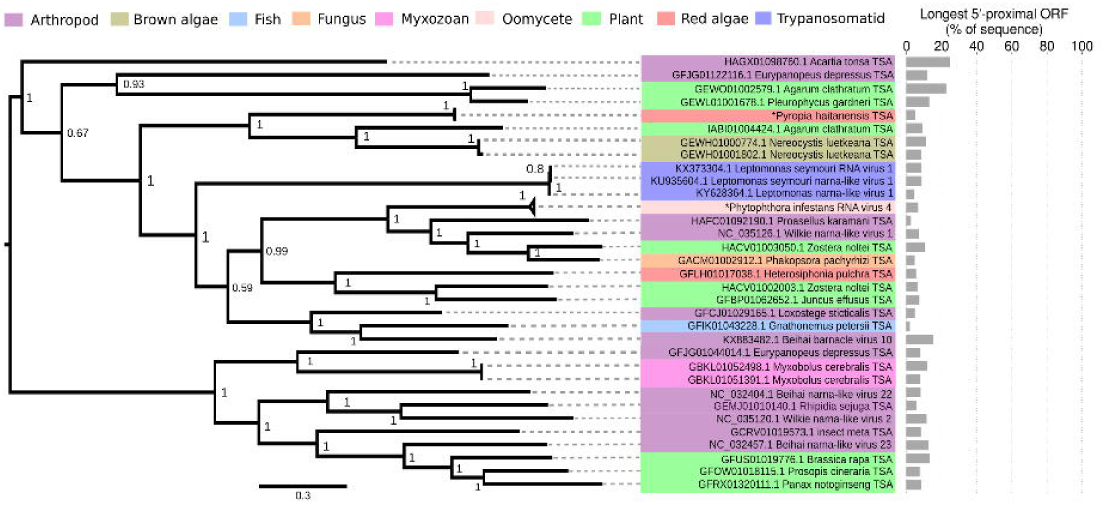
Midpoint-rooted Bayesian phylogenetic tree of betanarnaviruses. As in Figure 2, the longest 5′-proximal ORF in the negative-strand R0 frame is shown in the bars to the right. Five highly similar TSA sequences for *Pyropia haitanensis* and six highly similar sequences for Phytophthora infestans RNA virus 4 (both indicated with asterisks) are collapsed to single taxa. Accession numbers are for the nucleotide sequences from which the corresponding protein sequences were derived.

Notably, in both clades, a number of TSA sequences are highly divergent from other taxa in the phylogenies (**Figure 2** and **Figure 3**), suggesting that further sampling will continue to reveal new clades. Conversely, other taxa were disproportionately well sampled (e.g. nine Zhejiang mosquito virus 3 sequences with >95% pairwise amino acid identity were found; Shi et al. 2017).

### Genomic architecture and terminal regions

The terminal sequences of alphanarnaviral and betanarnaviral genomes were found to be dissimilar – the former group having short runs of G and C residues at the 5′ and 3′ termini, respectively, and the latter having A/U-rich termini with considerably longer 3′ untranslated regions (UTRs) (**Supplementary Figure S3**). Local RNA secondary structures are predicted to occur at the 5′ ends of the ONLV1 and ONLV2 genomes, coincident with a reduction in synonymous site variation in the RdRp ORF (see below). The putative structures are large (92 nt and 103 nt), and the G-rich 5′ terminus forms an integral component of the stem duplex (**Supplementary Figure S4A**), as it does in the *Saccharomyces* narnaviruses (Rodríguez-Cousiño et al. 1998; Fujimura and Esteban 2007). Meanwhile, the stop codons of the RdRp ORFs are predicted to be situated within shorter RNA stem-loop structures at the genomic 3′ termini (**Supplementary Figure S4, B and C**).

### Identification of ORFs in narnaviral genomes

We next determined for each sequence the longest stop codon-free region (ORF) in each of the three possible reading frames on each of the positive and negative strands, restricting to ORFs that begin within 200 nt of the 5′ end of the positive or negative strand, as appropriate (**Figure 4**). We used this approach because, under normal circumstances, non-5′-proximal ORFs are not expected to be translated, and the inclusion of spurious long non-5′-proximal ORFs might dilute the signal from translatable 5′-proximal ORFs. We designated the reading frame of the RdRp ORF as frame “F0” (for forward orientation, 0 frame), followed by frames “F+1” and “F+2” in the same orientation; whereas the reverse complement of the set of codons in frames F0, F+1 and F+2 were designated “R0” (reverse orientation, 0 frame), followed by “R+1” and “R+2” respectively.

**Figure 4.**
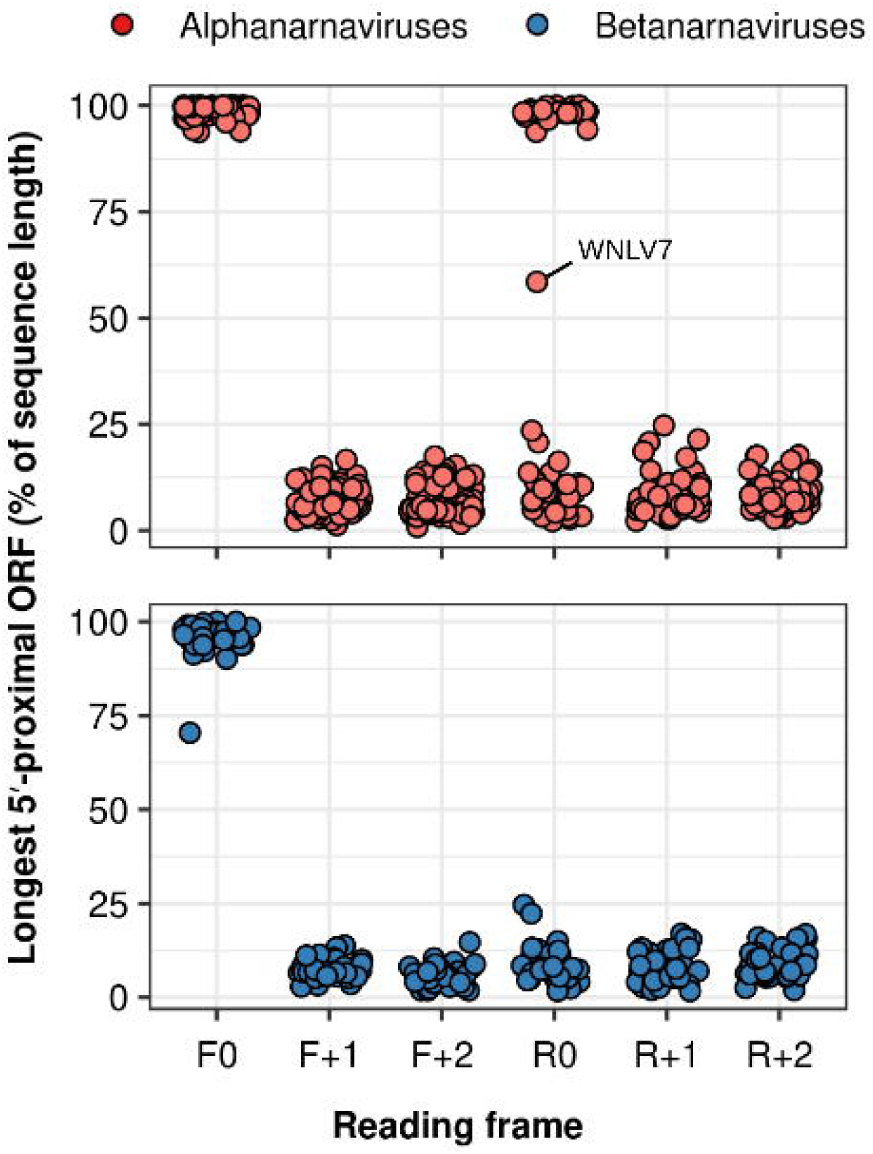
The longest 5′-proximal stop codon-free regions in each of the three possible positive strand and negative strand reading frames, for alphanarnaviruses (red) and betanarnaviruses (blue), as a percentage of the sequence length. Wenling narna-like virus 7 (WNLV7) has an intermediate-length R0-frame ORF, as indicated. Mean values are plotted for taxa with high levels of representation in the underlying data set (i.e. Zhejiang mosquito virus 3, Phytophthora infestans RNA virus 4, and TSA sequences from *Pyropia haitanensis*).

In line with the relatively short lengths of narnaviral untranslated regions (UTRs), the RdRp-encoding (i.e. frame F0) stop codon-free regions occupied 93.8–100.0% (median 99.5%) of alphanarnaviral sequences and 70.5–100.0% (median 97.0%) of betanarnaviral sequences (**Figure 4**). Several sequences in the data set are likely to be incomplete and, as a result, to have misannotated start or stop codons; for example, the genome of Hubei narna-like virus 18 (KX883517.1; Shi et al. 2016) is annotated as containing a 155-nt 5′ UTR, but this UTR contains no stop codons in-frame with the RdRp, and a blastx search of the sequence shows that it encodes amino acid sequence that shows close homology to the RdRp of the related Hubei narna-like virus 19 (e-value = 4 × 10^−12^).

Strikingly, other than the RdRp ORF, 5′-proximal stop-codon free regions occupying >25% of the genome were present only in the R0 frames and only in a subset of alphanarnaviral sequences, whereas stop (UAG, UAA, UGA) codons in all other frames were relatively common (**Figure 4**). The absence of stop codons in frame R0 in this subset directly mirrors an absence of the reverse-complementary codons (CUA, UUA, UCA) in the RdRp-encoding (F0) frame.

In a single case – Wenling narna-like virus 7 (GenBank: KX883602.1; Shi et al. 2016) – an rORF of intermediate length (531 codons; 58.5% of the sequence) was found (**Figure 4**), compared with an 890-codon stop-free RdRp-encoding region on the forward strand. The 5′ half of this RdRp ORF includes several codons corresponding to R0-frame stops: including two CUA (reverse-strand = UAG) codons, at positions 105 and 373; three UCA (reverse-strand = UGA) codons at positions 81, 134 and 157; and one UUA (reverse-strand = UAA), at position 72. However, the closely related sequences of Wenling narna-like virus 8 (KX883605.1), Beihai narna-like virus 24 (KX883500.1), and the TSA sequence from *Caridina multidentata* (IABX01132835.1) all contain rORFs occupying >90% of the respective sequences (**Figure 2**).

### Avoidance of CUA, UUA and UCA codons in RdRp-encoding ORFs

To enable a representative assessment of codon usage in narnaviral ORFs, sequences predicted to encode RdRp proteins with >90% amino acid identity were clustered using CD-HIT (Fu et al. 2012) and a single sequence from each cluster was retained for further analysis. After this step, 53 alphanarnavirus and 29 betanarnavirus representative sequences were left. An rORF occupying at least 90% of the sequence was present in 26 of the 53 alphanarnavirus sequences.

Among alphanarnaviruses, codon usage (as a proportion of total codons) in the RdRp ORF was broadly similar in taxa with and without long rORFs, with the exception of the three forward-orientation codons that introduce reverse-orientation stops (i.e. CUA, UUA and UCA) (**Figure 5**). Effectively by definition, each of these three codons is excluded from large portions of sequences with long rORFs (mean usage per associated amino acid in RdRp ORF = 0.0037, 0.0054 and 0.0056, respectively), but are 17- to 26-fold more common in those alphanarnaviruses without long rORFs (mean usage per associated amino acid = 0.0973, 0.0896, and 0.1277, respectively) (**Figure 6A**). These three codons were also relatively common in the RdRp ORFs of the betanarnaviruses (mean usage per associated amino acid = 0.1323, 0.1452 and 0.1784, respectively). CUA and UUA both encode leucine (Leu) whereas UCA encodes serine (Ser). Examination of codon usage bias for these two amino acids shows that sequences with long rORFs specifically avoid these codons and use proportionately more of each alternative codon to encode these two amino acids (**Figure 6A**).

**Figure 5.**
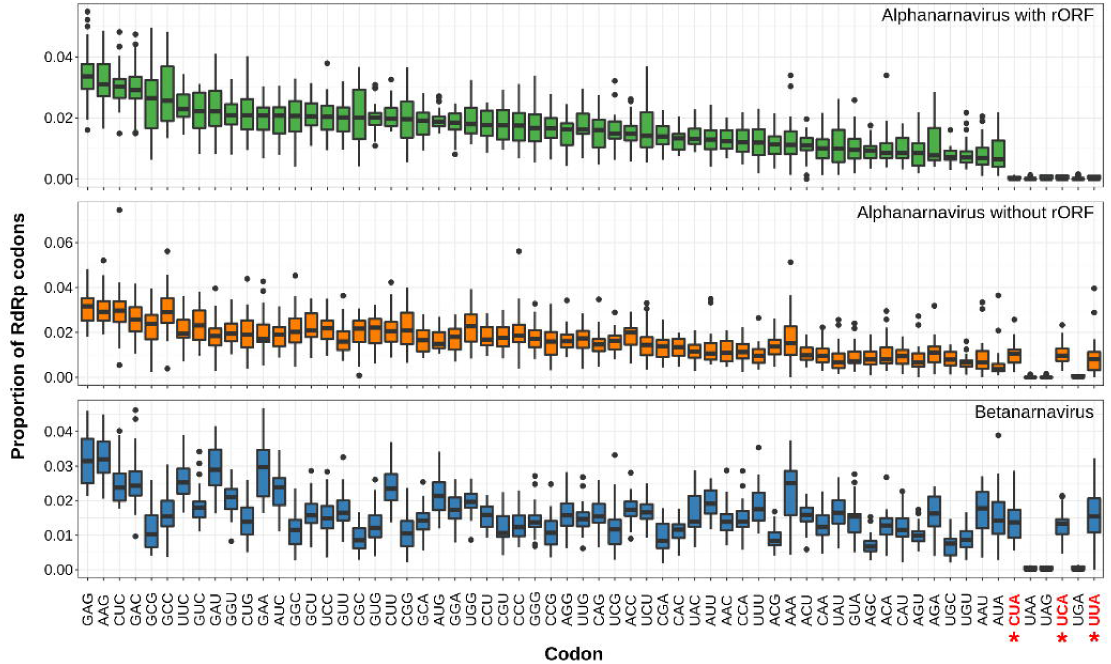
Box plots of codon usage (proportion of total codons) in RdRp ORFs. The upper and lower hinges are located at the first and third quartiles, respectively; the median values are indicated as horizontal lines; and the whiskers extend from the hinges to the furthest data points within 1.5 times the interquartile range (IQR) of the hinges. Data more than 1.5 times the IQR from the hinges are drawn as individual points (outliers). Codons are ordered according to their median frequency in alphanarnaviruses that contain the rORF. Alphanarnaviruses that contain the rORF show a specific avoidance of the three codons marked with red asterisks (CUA, UUA and UCA), which correspond to the reverse complements of stop codons.

**Figure 6.**
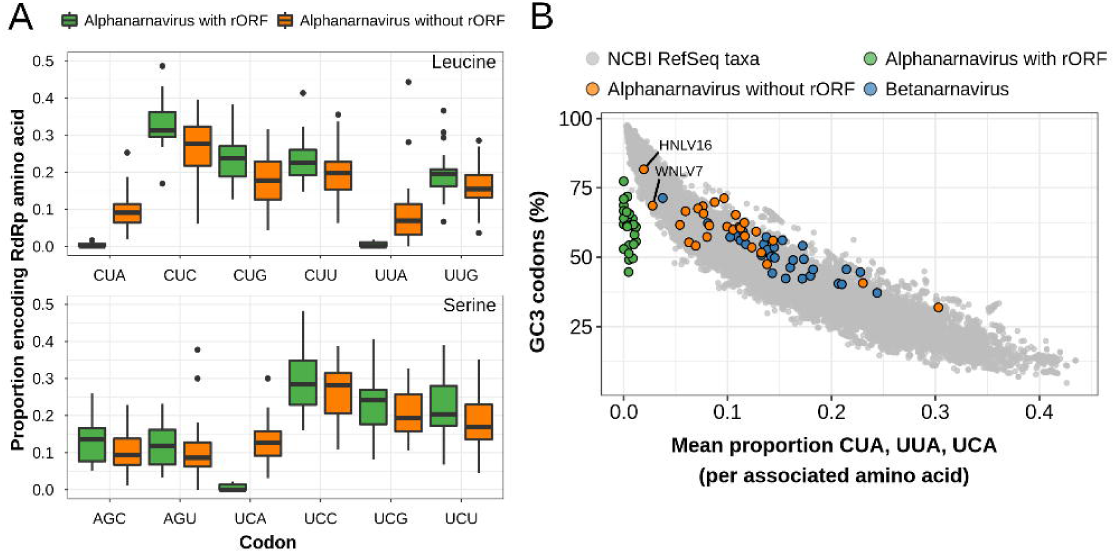
Specific selection against CUA, UUA and UCA codons in rORF-containing alphanarnaviruses. **(A)** Box plots of codon usage bias (proportion of codons encoding the amino acid) for leucine and serine. Box plots are drawn as in Figure 5. **(B)** Comparison of mean usage of CUA, UUA and UCA (per associated amino acid) with the third-position GC (GC3) content of codons. Prokaryotic and eukaryotic taxa in NCBI RefSeq are shown as grey points. The RdRp ORFs of non-rORF alphanarnaviruses, rORF alphanarnaviruses and betanarnaviruses are indicated with orange, green and blue points, respectively. The Wenling narna-like virus 7 (WNLV7) and Hubei narna-like virus 16 (HNLV16) sequences are indicated.

### Comparison of codon usage across species

Like all (+)ssRNA viruses, narnaviruses are dependent upon host tRNA pools and translation machinery for their gene expression. This raises the possibility that the paucity of CUA, UUA and UCA codons in the rORF-containing sequences could at least partially reflect an adaptation to particular host codon usage patterns.

Using the RefSeq database from NCBI, as tabulated in the latest release of the codon usage table database (CUTD) (Athey et al. 2017), we assessed the relative proportions of the different leucine and serine codons across species. We found that global usage of CUA, UUA and UCA (per associated amino acid) scales directly and inversely with GC content at the third position of codons (GC3) and only species with extremely high GC3 had bias against CUA, UUA and UCA as extreme as that in the rORF-containing narnaviruses (**Figure 6B**). Most taxa with mean overall CUA, UUA and UCA usage as low as that of rORF-containing narnaviruses are very high-GC bacteria and archaea (mean GC3 content = 95.2%). The eukaryotic taxon with lowest mean overall usage of these three codons (0.0107, cf. mean for rORF-containing narnaviral sequences = 0.0049) was a fungus, *Rhodotorula graminis* (phylum Basidiomycota), which has a GC3 content of 90.4%. In contrast, the mean GC3 contents of rORF-containing and non-rORF-containing alphanarnaviral sequences were considerably lower (61.1% and 59.9%, respectively; **Figure 6B**). Thus, selection against CUA, UUA and UCA codons in rORF-containing alphanarnaviruses cannot readily be explained by GC bias or host codon usage bias.

To investigate whether codon usage could be used as an indicator of the likely host taxonomic group or groups for narnaviruses, and motivated by previous work (Kapoor et al. 2010), we performed a principal component analysis (PCA) of codon usage statistics for host groups that frequently co-occur with narnaviruses in transcriptomic datasets. The PCA allowed clear segregation of arthropods from ascomycetes and basidiomycetes (two major fungal phyla) and streptophytes (land plants and some green algae), but the latter three groups did not clearly segregate from each other (**Supplementary Figure S5**). Narnaviral codon usage statistics were projected onto the resulting principal component space, and did not clearly segregate with a single host group, regardless of whether or not leucine and serine codons were included in the analysis (**Supplementary Figures S5 and S6**). Chordates were excluded from this analysis, as their codon bias is heavily influenced by the avoidance of CpG-ending codons, and they group distinctly from other phyla and from the narnaviruses.

### Analysis of amino acid and nucleotide conservation in the rORF

To assess conservation within the rORF of the representative narnavirus sequences, the predicted RdRp amino acid sequences were aligned and then back-translated to RNA. Overall, the amino acid sequences predicted to be encoded by rORFs were rather more divergent than the corresponding set of RdRps (mean pairwise identities 20.1% and 26.6%, respectively). Codon-based alignments of RdRp ORFs and rORFs were generated, based in both cases on the RdRp amino acid alignments, and amino acid conservation and synonymous site variation of codons (Firth 2014) were assessed (**Figure 7**).

**Figure 7.**
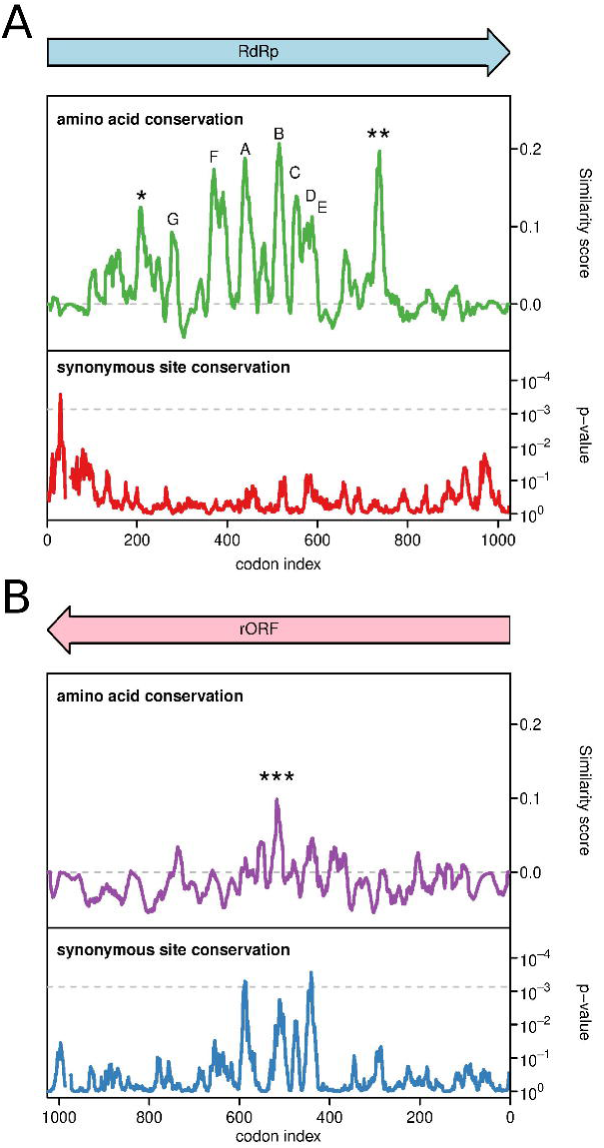
Amino acid conservation and synonymous site conservation in rORF-containing alphanarnaviruses. (**A**) RdRp amino acid conservation, and the corresponding synonymous site conservation analysis. In the latter, the grey dashed line indicate an approximate 5% false positive threshold after correcting for multiple tests (i.e. ~69 × 15-codon windows in the 1027-codon ORF). (**B**) The equivalent plots for the rORF. In each case, conservation was assessed in a 15 codon/amino acid window. Motifs A–G in the RdRp amino acid sequence are indicated with letters. Additional consensus motifs indicated with asterisks are as follows: * = R&UP, ** = Pxx[L/V]GGx[G/N]xP and *** = QvxExExxPREREAH; U = I, L, V or M, & = I, L, V, M, A, P, G, F, W or Y.

The core RdRp functional motifs (A to G; see above) localise to a region of fewer than 400 amino acids; with additional short conserved motifs (R&UP and Pxx[L/V]GGx[G/N]xP; U = I, L, V or M, & = I, L, V, M, A, P, G, F, W or Y) being found further upstream and downstream, respectively (**Figure 7A**). Increases in synonymous site conservation were observed in the rORF directly opposite RdRp motifs A and E, and – to a lesser extent – motifs B, C, D and F, and at the Pxx[L/V]GGx[G/N]xP motif (**Figure 7B**) indicating that these conserved motifs in the RdRp constrain synonymous site variation in the rORF.

The largest increase in synonymous site conservation in the RdRp ORF lies between codons 23 and 37 (**Figure 7A**), coinciding with the location of a known RdRp interaction site and a *cis*-acting replication signal in the ScNV-20S genome (Fujimura and Esteban 2007). This region also contains the predicted 5′ structures shown in **Supplementary Figure S4A**. Increased synonymous site conservation was independently observed in this region for betanarnaviruses (not shown). The most highly conserved region in the putative rORF-encoded amino acid sequence was also directly opposite the RdRp motif B, in a 15-amino acid window of the alignment centred on position 510 (**Figure 8B**), where the rORF-encoded consensus sequence is QvxExExxPREREAH.

## DISCUSSION

In this study, we have shown that narnaviral sequences form two major clades, which we suggest to be formally recognized as a pair of genera. We show that sequences with long rORFs are restricted to the proposed genus *Alphanarnavirus*, which also contains the prototypical *Saccharomyces cerevisiae* 20S and 23S narnaviruses. The proposed genus *Betanarnavirus* contains pathogens of diverse unicellular eukaryotes, including the oomycete *Phytophthora infestans* and the trypanosomatid *Leptomonas seymouri*. The two proposed genera differ in terms of their relative codon usage preferences and in the composition of their genomic termini, with alphanarnaviruses having GC-rich termini and a higher overall GC3 content in codons, and the betanarnaviruses having AU-rich termini. These variations might, to some extent, reflect divergence in host taxa. Although some of the available sequences are likely incomplete, there is evidence that members of both proposed genera exhibit complementarity between the genomic termini. In ScNV-20S and ScNV-23S, the 5′-GGGG and CCCC-3′ genomic termini, besides 5′ and 3′ RNA structures, are essential for efficient virus replication (Esteban and Fujimura 2003; Fujimura and Esteban 2004; Esteban et al. 2005). Since narnavirus RNAs are non-polyadenylated and probably non-capped, they are vulnerable to host mRNA degradation pathways. However, the viral RdRp binds to the 3′-terminal CCCC and an adjacent RNA stem-loop structure in the positive strand and this is thought to stabilize the genome and protect the 3′ end from host 3′ exonuclease degradation (Fujimura and Esteban 2004; Fujimura and Esteban 2007). Similar elements also exist at the 3′ end of the negative strand (Esteban et al. 2005; Fujimura and Esteban 2007). Meanwhile, the positive-strand 5′-proximal RNA structure protects the genome from the host SKI1/XRN1 5′ exonuclease (Esteban et al. 2008). These stability elements are thought to be particularly important for narnaviruses due to the lack of capsids or membrane-associated replication that other (+)ssRNA viruses use to protect their genomes (Fujimura and Esteban 2007).

The diversity of sources of co-clustering narnaviral genomes suggests either that horizontal transfer of narnaviruses between divergent hosts has occurred, or that at least some of these sources are not the *bona fide* host species, potentially reflecting contamination from parasitic or commensal organisms, gut contents, or external debris. For example, in **Figure 2**: two TSA sequences from the fly-infecting fungus *Entomophthora muscae* (GENC01006608.1 and GEND01011317.1) cluster within a clade of arthropod-derived sequences; a TSA sequence (GFKT011160020.1) from the spider *Nephila clavipes* has 95% nucleotide identity to ScNV-20S (AF039063.1); and two TSA sequences (GGCO01105932.1 and GGCO01034162.1) from the barley plant *Hordeum vulgare* cluster with sequences derived from the Basidiomycota obligate plant pathogenic fungi *Uromyces appendiculatus* and *Puccinia striiformis*. Since ScNV-20S is a well-characterized virus of *Saccharomyces cerevisiae*, it is likely that the *Nephila clavipes* sequence derives from fungal contamination. The *Entomophthora muscae* sequences were obtained from fungus-infected *Delia radicum* cabbage flies, and indeed the NCBI SRA taxonomy analysis webpages (alpha version, 30 May 2019) for the corresponding RNA-Seq libraries show 4–16 times as much fly RNA as fungal RNA (within the only 8–9% of reads that were taxonomically identified); thus these TSAs may derive from infected insect cells. Similarly, the *Hordeum vulgare* TSAs may derive from infected fungal cells since the NCBI SRA taxonomy analysis webpages show contamination with Opisthokonta (i.e. animal/fungi) RNA at ~4% the level of plant RNA. Thus, it is important to apply caution when attempting to infer the host of TSA sequences.

Given the co-occurrence of fungal sequences in arthropod transcriptomic data sets, it was originally suggested that the identified narnavirus sequences might derive from fungal contaminants (Chandler et al. 2015; Cook et al. 2013). More recently, evidence was put forward that at least some narnaviruses are true arthropod viruses: for example, the group of alphanarnaviruses comprising KF298275.1, KF298276.1, KF298284.1, Zhejiang mosquito virus 3, KP642119.1 and KP642120.1 (**Figure 2**) come from four different studies and five different Culicinae mosquito species (Chandler et al. 2015; Cook et al. 2013; Shi et al. 2016; Shi et al. 2017); moreover, in several samples, viral RNA accounts for >0.1% (in one case, >2%) of total non-ribosomal RNA reads, which appears unlikely if the virus is infecting a contaminant (Shi et al. 2016; Shi et al. 2017). Very recently, another narnavirus in this clade (MK628543.1; Culex narnavirus 1; 97% nt identity to KP642120.1) was found to persistently infect a *Culex tarsalis* cell culture and also give rise to typical 21-nt viral siRNAs with equal coverage of both strands, indicative of active infection (Göertz et al. 2019).

An ONLV2-related endogenized virus element (EVE) has also been reported for arthropods (Shi et al. 2016); EVEs represent virus fragments that have been spuriously reverse-transcribed and integrated into host genomes, and provide further evidence as to the true host(s) of virus groups (Katzourakis and Gifford 2010). We also detected sequences predicted to encode proteins with significant homology to the RdRp proteins of rORF-containing narnaviruses (tblastn e-values ≤ 1.0 10^−10^) in whole genome shotgun (WGS) assembly data sets from several arthropods, including several species of ant (*Aphaenogaster floridana, Aphaenogaster rudis, Lasius niger, Linepithema humile* and *Cataglyphis niger*), the seven spot ladybird (*Coccinella septempunctata*) and the glassy-winged sharpshooter (*Homalodisca vitripennis*). Thus there is now strong evidence that at least some rORF-containing alphanarnaviruses are *bona fide* arthropod viruses. On the other hand, other rORF-containing alphanarnaviruses come from fungal (or plant) samples that have no obvious association with arthropods (< 0.8% metazoan-mapping reads in total per sample), such as the *Uromyces appendiculatus* and *Puccinia striiformis* samples, indicating that presence of the rORF is likely not a unique adaptation to arthropod infection.

Since our original identification of long rORFs in the ONLV1 and ONLV2 genomes, besides sequences from *Uromyces appendiculatus* and *Puccinia striiformis* transcriptomes (Cook et al. 2013), a number of other studies have discovered narnaviral genomes containing similar rORFs (Chandler et al. 2015; Shi et al. 2016; Shi et al. 2017; Viljakainen et al. 2018; Göertz et al. 2019). These rORF-containing sequences appear not to form a monophyletic group, although they do show considerable phylogenetic clustering. The rORF occurs exclusively in the R0 frame relative to the RdRp ORF, and large sections of the latter therefore exhibit a specific exclusion of the three codons CUA, UUA and UCA, which are however widely used in the RdRp ORF of non-rORF-containing narnaviruses. Nonetheless, these codons *are* still used occasionally in rORF-containing narnaviruses, exclusively within the 5′-most extremity (3%) of the RdRp coding region (at or upstream of the rORF stop codon). This indicates that selection against these codons is not due to an inability for these codons to be decoded by the host translational machinery (e.g. due to the lack of cognate tRNAs). This is also supported by our analysis of cellular organism codon usage which showed that the vast majority of organisms with GC3 in the same range as rORF-containing narnaviruses have CUA, UUA and UCA mean codon usage values well above those of the rORF-containing narnaviruses (**Figure 6B**). Thus, the presence of the long rORF in highly divergent alphanarnaviruses cannot be explained as an artefact of RdRp ORF codon usage or GC bias.

To our knowledge there is no experimental evidence for protein-coding ORFs in the negative strand of any (+)ssRNA virus. A recent bioinformatic study used single-sequence randomization procedures to identify ORFs overlapping previously annotated ORFs in RNA virus sequences that are statistically significantly longer than expected by chance, with the assumption being that such ORFs are likely to be functional (Schlub et al. 2018). The authors identified statistically significantly long negative-strand stop-codon-free regions overlapping positive-strand ORFs in sweet potato virus 2 (NC_017970.1; family *Potyviridae*), Macrophomina phaseolina tobamo-like virus (NC_025674.1; family *Virgaviridae*), Nhumirim virus (NC_024017.1; family *Flaviviridae*), hibiscus chlorotic ringspot virus (NC_003608.1; family *Tombusviridae*), hydrangea ringspot virus (NC_006943.1; family *Alphaflexiviridae*), and Scrophularia mottle virus (NC_011537.1; family *Tymoviridae*). However it remains unclear how these ORFs would be translated. In eukaryotes, translation normally relies on recruitment of pre-initiation ribosomes to the 5′ end of mRNAs, followed by scanning and initiation at the first AUG in a good initiation context. Thus, the majority of *de novo* protein-coding ORFs tend to evolve towards the 5′ ends of transcripts, frequently overlapping the 5′ end of an ancestral protein-coding ORF where they can be translated via a process known as leaky scanning (Firth and Brierley 2012). If protein-coding ORFs exist in the negative strand of (+)ssRNA viruses, one might expect the majority to have initiation sites close to the 5′ end of the negative strand. However for the six ORFs above there are, respectively, 171, 101, 115, 8, 3 and 100 intervening AUG codons between the 5′ end of the negative strand and the first in-frame AUG codon in the ORF, which would appear to rule out 5′-end-dependent scanning as a translation mechanism. 5′-distal ORFs on positive-sense transcripts are often translated via ribosomal frameshifting or stop codon readthrough, but that still requires 5′-proximal initiation in the pre-frameshift/pre-readthrough ORF (Firth and Brierley 2012). 5′-distal ORFs are occasionally translated via an internal ribosome entry site (IRES), but IRESes are normally complex elements and it would be difficult for them to evolve within protein-coding sequences. Simple inefficient IRESes are a possibility and, compounded with the low availability of negative strand, would lead to extremely low expression levels of any resulting proteins. Splicing is unknown in (+)ssRNA viruses as they all replicate cytoplasmically. Subgenome-sized negative-sense transcripts, where they are produced, are normally 5′ co-terminal with the full-length negative strand so would not normally provide access to internal ORFs, although the production of other classes of negative-sense transcripts is plausible (Sztuba-Solińska et al. 2011). Given the pronounced evolutionary adaptations that might be required to express such negative-strand ORFs, in the absence of experimental data they would be more plausible if they were conserved in related species. However this has not been demonstrated. For example, the Nhumirim virus rORF is not conserved in the next most closely related flavivirus genome sequence, that of Barkedji virus (MG214906.1; 71% aa identity in the forward ORF) where the rORF region is disrupted by eight stop codons. Similarly, the sweet potato virus 2 rORF is not conserved in sweet potato virus G (NC_018093.1; 74% aa identity in the forward ORF) where the rORF region is disrupted by nine stop codons. Indeed it is not even conserved in other isolates of sweet potato virus 2 – e.g. KP729268.1 and KP115618.1 each have a stop codon. Thus we contend that the alphanarnavirus rORF is currently the only plausible candidate for a negative-strand coding ORF in (+)ssRNA viruses.

The replication of most (+)ssRNA viruses is membrane-associated (den Boon and Ahlquist 2010; Romero-Brey and Bartenschlager 2014; Shulla and Randall 2016; Ertel et al. 2017). Infection typically results in extensive rearrangement of host membranes to produce membrane-bound mini-organelles within which genome replication occurs. These structures co-localize viral RNA and proteins and may also shield viral positive:negative strand duplexes from recognition by host dsRNA-recognizing antiviral proteins such as RIG-I, MDA-5, PKR and the RNAi machinery. Sequestering replication within membranous compartments may also play a role in separation of translation from replication: positive-sense mRNAs are extruded into the cytoplasm for translation, whereas the negative strand remains protected within the membranous compartment for replication. This prevents ribosomes translating 5′ to 3′ from colliding with RdRps tracking 3′ to 5′ on the same template. Thus, at least at later time points, the negative strand is expected to be generally sequestered away from the translational machinery. Further, the negative strand of (+)ssRNA viruses often lacks the 5′ structures (such as a 5′ cap, 5′ covalently linked viral protein of the genome, or IRES structure) required to efficiently recruit ribosomes. Narnavirus replication is, however, quite atypical among eukaryote-infecting (+)ssRNA viruses, and this may explain why they – possibly uniquely – appear to have evolved negative-strand coding capacity. Narnavirus replication is thought to occur entirely within the cytoplasm, not associated with cellular membranes (Solórzano et al. 2000; Fujimura et al. 2005). As mentioned above, the narnavirus RdRp is more closely related to the RdRp of bacteriophages in the family *Leviviridae* than it is to the RdRp of other (+)ssRNA viruses. Similar to leviviruses, the RNA in narnavirus replication intermediates is essentially single-stranded (Blumenthal and Carmichael 1979; Dobkin et al. 1979; Zinder 1980; Takeshita and Tomita 2012; Fujimura et al. 2005; Wickner et al. 2013). Each newly synthesized RNA is released before a new round of replication commences and resting complexes comprising a negative-sense or positive-sense single-stranded RNA bound to a single copy of the viral RdRp are present within the cytoplasm (Solórzano et al. 2000). Thus, for narnaviruses, the positive and negative strands likely have similar accessibility to the host cell translational machinery.

The rORF is present in many but not all alphanarnaviruses. Surprisingly, the distribution of the rORF does not appear to be monophyletic. Thus it may have evolved multiple times, or it may have been present ancestrally and lost from some lineages. The observation that the rORF is always in the R0 frame relative to the RdRp ORF may be evidence against independent evolution in multiple lineages, although it could also be a consequence of codon bias, e.g. if long ORFs are by chance more likely to occur in the R0 frame as a result of RdRp ORF codon usage. Such random ORFs could provide “seeds” for the evolution of longer overlapping genes (Belshaw et al. 2007). Another surprising feature is that the rORF, where present, is nearly always full-length. If the sole function of the rORF were to encode an additional protein, one might expect to see a variety of rORF lengths in different lineages whereas we only see a single example of an rORF beginning within 200 nt of the 5′ end of the negative strand with length in the range 24–94% of the full sequence length (**Figure 4**). We found scant evidence for amino acid conservation in the rORF. It is possible that the very high divergence between most of our sequences might obscure a weak conservation signature (in contrast to the RdRp which is the most highly conserved RNA virus protein known). However it is also possible that the protein product of the rORF is not itself functionally important. A possible alternative explanation is that rORF translation in itself might facilitate replication e.g. by disassociating dsRNA, or by increasing negative strand RNA stability (Bicknell and Ricci 2017). Alternatively, if the negative strand is unavoidably accessible for translation, exon-junction-complex-independent nonsense mediated decay may provide strong selection pressure against non-full-length rORFs (Kurosaki et al. 2019).

Although, as discussed above, the negative strands of most (+)ssRNA viruses are normally expected to be occluded in membranous compartments, it is currently unclear whether at early timepoints – before extensive membranous compartments have formed – negative strands produced in a first round of replication might be exposed to the cytoplasm and potentially available for translation (Shulla and Randall 2015). It is also not inconceivable that some (+)ssRNA viruses might have evolved mechanisms to allow some negative strands access to the translational machinery even at late timepoints. If so, it is possible that some (+)ssRNA viruses might have evolved ribosome recruitment elements within their negative strands. Thus it is perhaps too early to dismiss the possibility of negative-strand ORFs in other (+)ssRNA virus lineages. In most cases, however, any proteins encoded on the negative strand would be expected to be expressed at much lower levels than positive-strand encoded proteins simply because of the huge disparity in positive:negative RNA abundance during virus infection (typically of order 100:1; Novak and Kirkegaard 1991; Kopek et al. 2007; Irigoyen et al. 2016). Even for narnaviruses, less than 1–2% of ScNV-20S viral RNA in infected cells is negative-sense (Rodriguez-Cousiño et al. 1991; Fujimura et al. 2005), indicating that the rORF, where present, is likely to be expressed at a much lower level than the RdRp.

In conclusion, we have provided the first strong evolutionary evidence for reverse coding capacity in a group of positive-sense RNA viruses. The alphanarnavirus rORFs are exceedingly long compared to most known overlapping genes (Pavesi et al. 2018; Brandes and Linial 2016). Thus their study is not only of interest to the evolution and molecular biology of viruses, but also of broad relevance to understanding the evolution of overlapping genes and the *de novo* origin of genes.

## METHODS

The NCBI non-redundant (nr) and transcriptome shotgun assembly (TSA) databases were downloaded on August 31, 2018. Narnaviral sequences were identified in these databases using, tblastn (version 2.8.0) (Camacho et al. 2009), with the RdRp protein sequence of ScNV-20S and the updated RdRp sequence of ONLV1 (completed in this study) as queries. In total, 81 sequences were found by combined searches using these queries (46 in the nr database and 35 in the TSA database). Hidden Markov model (HMM)-based searches were carried out using hhsearch (Finn et al. 2011), based on an alignment of RdRps from rORF-containing narnaviruses, leading to the identification of a further 43 TSA database sequences.

Phylogenetic trees were generated with MrBayes (version 3.2.7) (Huelsenbeck and Ronquist 2001), using a mixed substitution model with sampling across fixed amino acid rate matrices (aamodelpr = mixed) and 5,000,000 generations. All other parameters were set as defaults.

RNA structures in the genomic terminal regions were predicted by scanning full-length narnaviral genomes with RNALfold (version 2.4.9) (Lorenz et al. 2011), allowing a maximum base-pair span of 150 nt.

For all comparisons of alphanarnaviruses “with” and “without” rORFs, taxa in which the longest stop codon-free region (beginning within the 5′-most 200 nt) occupied at least 90% of the genomic sequence were assigned to the former group, whereas other taxa were assigned to the latter group.

A codon-based multiple sequence alignment of RdRp ORFs was produced by aligning the RdRp amino acid sequences with MUSCLE (version 3.8.31) (Edgar 2004) and then backtranslating to a nucleotide sequence alignment using the tranalign program in the EMBOSS suite (version 6.6.0.0) (Rice et al. 2000). The alignment was then mapped to ONLV2 sequence coordinates by removing alignment positions that contained a gap character in the ONLV2 sequence. Synonymous site variation was assessed using synplot2 (Firth 2014), with a window size of 15 codons. Comparison of amino acid sequences was performed using plotcon from the EMBOSS suite (version 6.6.0.0) (Rice et al. 2000) with a BLOSUM62 substitution matrix and a window size of 15 amino acids. When analysing synonymous site usage in the rORF, the reverse complement of the RdRp alignment was used, and aligned nucleotides on the reverse strand were converted to the corresponding protein sequences using AMAS (Borowiec 2016).

Principal component analysis (PCA) of codon usage across species was performed using relative codon abundances (i.e. normalized by the abundances of the associated amino acids). The principal components were calculated using the RefSeq sequences of cellular organisms, and narnaviral codon abundances were projected onto the resulting principal component space.

## Supporting information

Supplementary Data Set S1

## ACKNOWLEDGEMENTS

We thank Chris McCormick, Katherine Brown, Valeria Lulla and Hazel Stewart for useful discussions. Work was supported by Wellcome Trust grant [106207] and European Research Council grant [646891] to A.E.F.

