## Supplementary Figures for "A case for a reverse-frame coding sequence in a group of positive-sense RNA viruses"

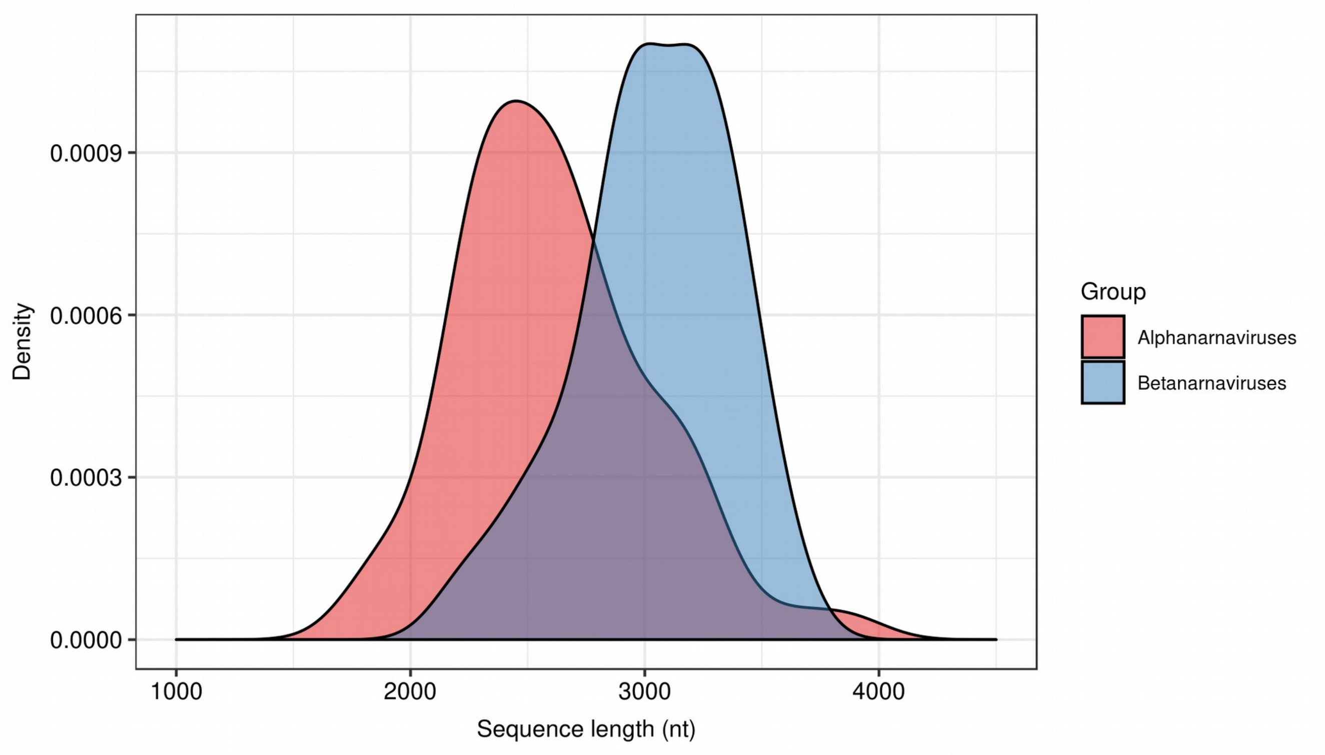


**Figure S1**. Sequence length distributions for alphanarnaviruses (*n* = 64) and betanarnaviruses (*n* = 33) included in the analysis.

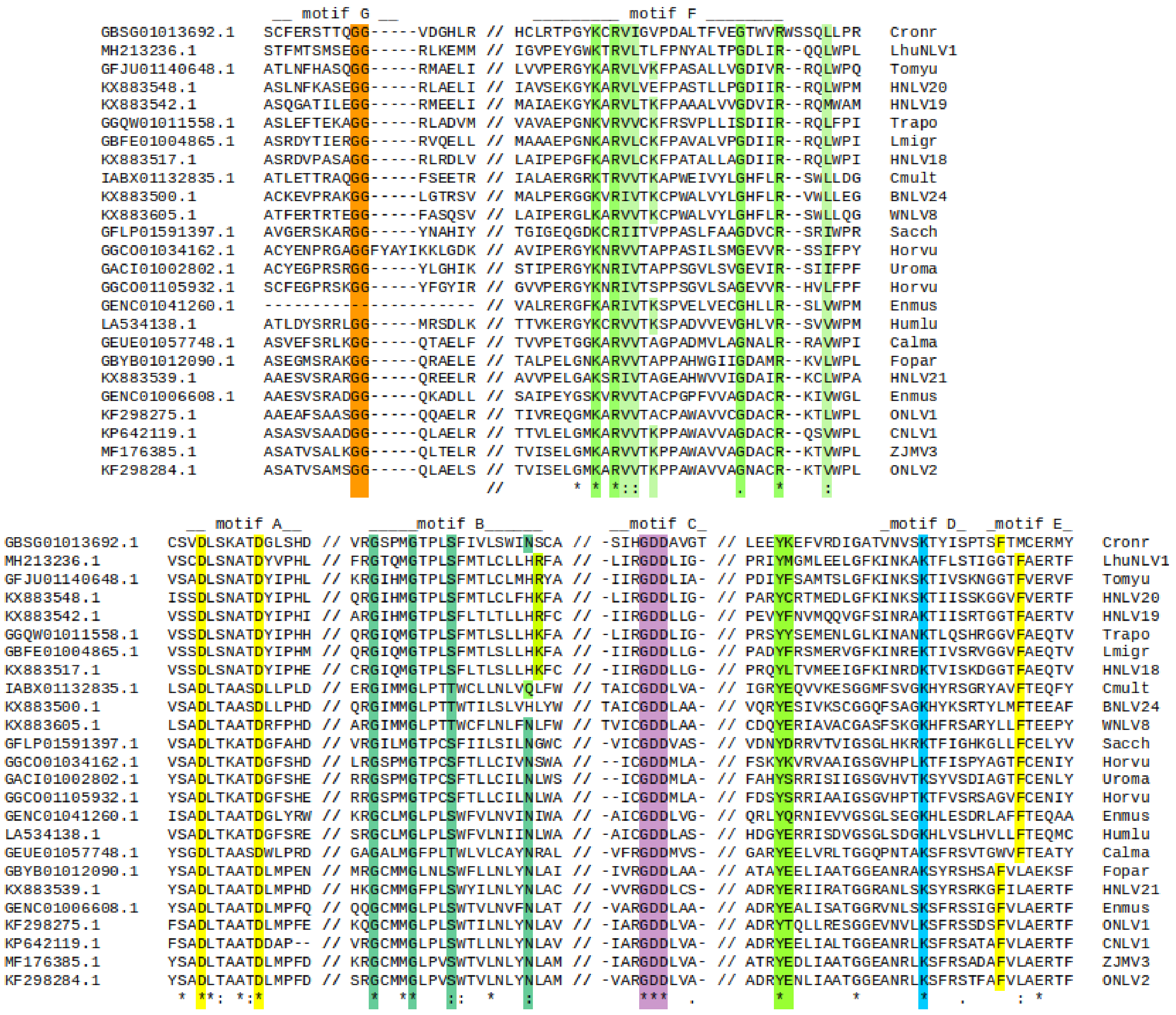


**Figure S2.** Alignment of RdRp protein-coding sequences of rORF-containing alphanarnaviruses. Known RdRp functional motifs are indicated and key functional residues previously described are highlighted. Key: Cronr = *Cronartium ribicola* TSA; LhuNLV1 = Linepithema humile narna-like virus 1; Tomyu = *Tomicus yunnanensis* TSA; HNLV = Hubei narna-like virus; Trapo = *Tracheliastes polycolpus* TSA; Lmigr = *Locusta migratoria* TSA; BNLV = Beihai narna-like virus; Cmult = *Caridina multidentata* TSA; WNLV = Wenling narna-like virus; Sacch = *Saccharum* hybrid TSA; Horvu = *Hordeum vulgare* TSA; Uroma = *Uromyces appendiculatus* TSA; Enmus = *Entomophthora muscae* TSA; Humlu = *Humulus lupulus* TSA; Calma = *Callosobruchus maculatus* TSA; Fopar = *Fopius arisanus* TSA; ONLV = Ochlerotatus-associated narna-like virus; CNLV = Culex-associated narna-like virus; ZJMV3 = Zhejiang mosquito virus 3.

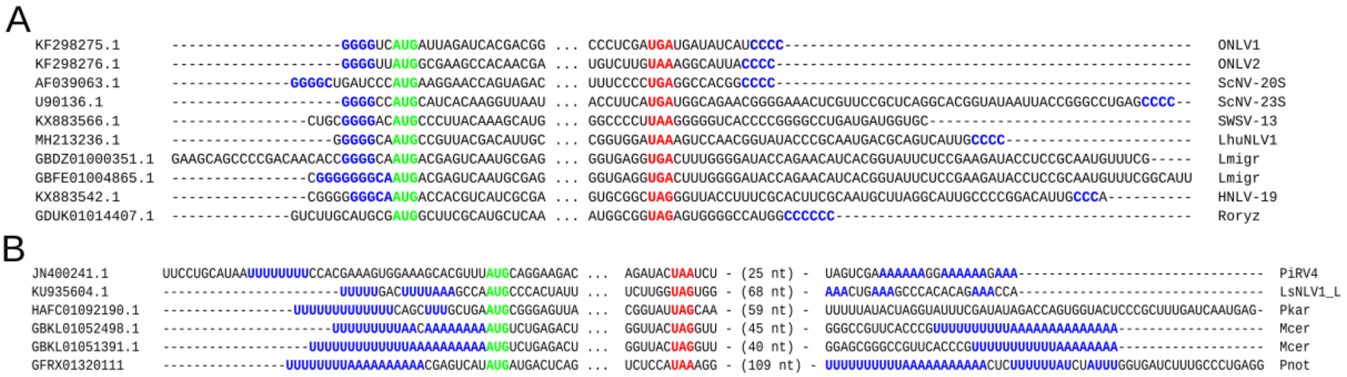


**Figure S3**. Alignments of **(A)** alphanarnaviral and **(B)** betanarnaviral genomic termini. The start and stop codons of the RdRp ORFs are shown in green and red, respectively. Complementary G/C- and A/U-rich regions are indicated in blue. Key: ONLV = Ochlerotatus-associated narna-like virus; ScNV-20S = Saccharomyces cerevisiae 20S narnavirus; ScNV-23S = Saccharomyces cerevisiae 23S narnavirus; SWSV-13 = Sanxia water strider virus strain 13; LhuNLV1 = Linepithema humile narna-like virus 1; Lmigr = *Locusta migratoria* TSA; HNLV-19 = Hubei narna-like virus 19; Roryz = *Rhizopus oryzae* TSA; PiRV4 = Phytophthora infestans RNA virus 4; LsNLV1_L = Leptomonas seymouri narna-like virus 1 segment L; Pkar = *Proasellus karamani* TSA; Mcer = *Myxobolus cerebralis* TSA; Pnot = *Panax notoginseng* TSA.

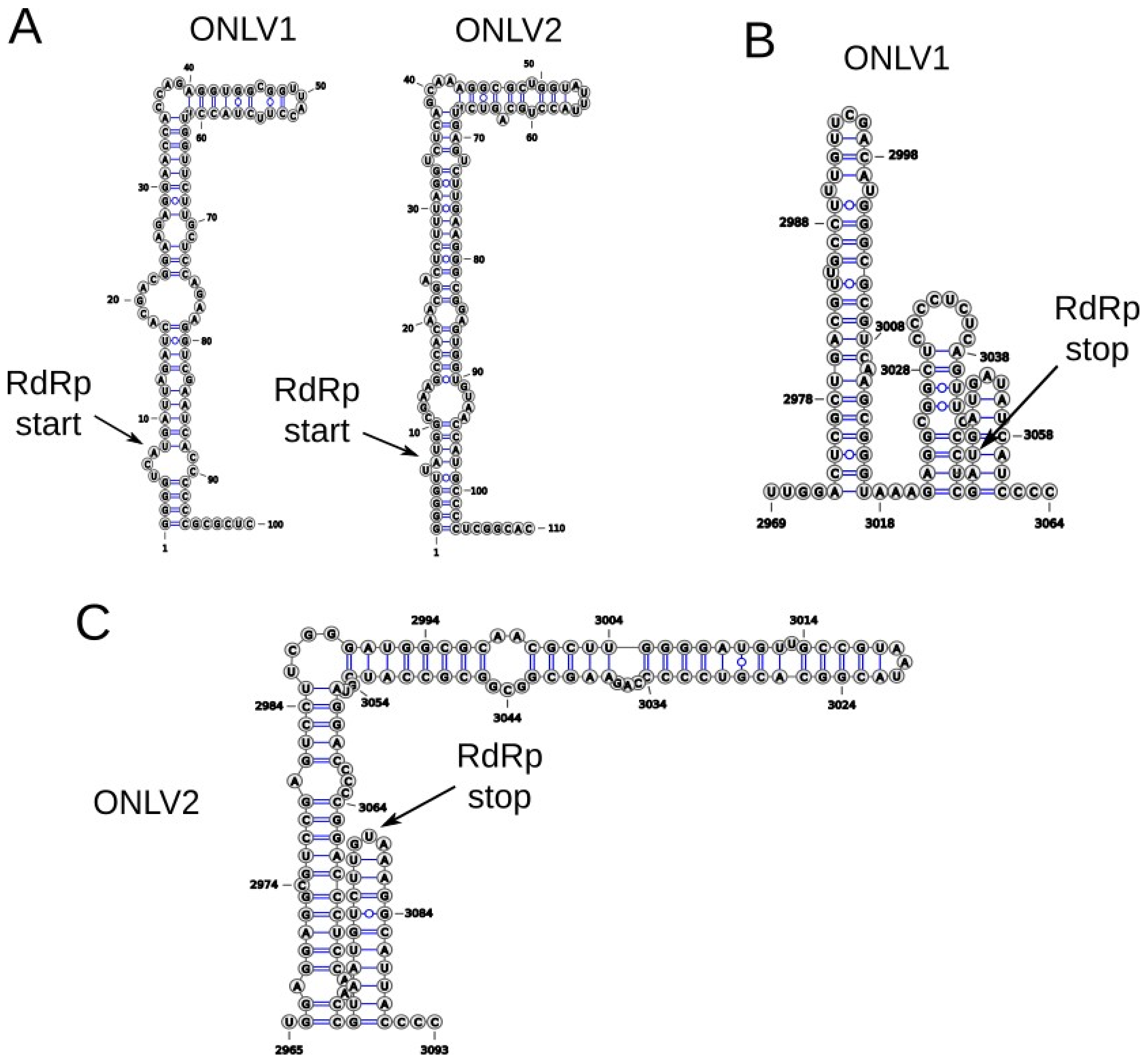


**Figure S4**. Predicted RNA secondary structures at the terminal regions of alphanarnaviral genomes. **(A)** 5′ termini of *Ochlerotatus*-associated narna-like virus (ONLV) 1 and ONLV2; **(B)** 3′ terminus of ONLV1; and **(C)** 3′ terminus of ONLV2.

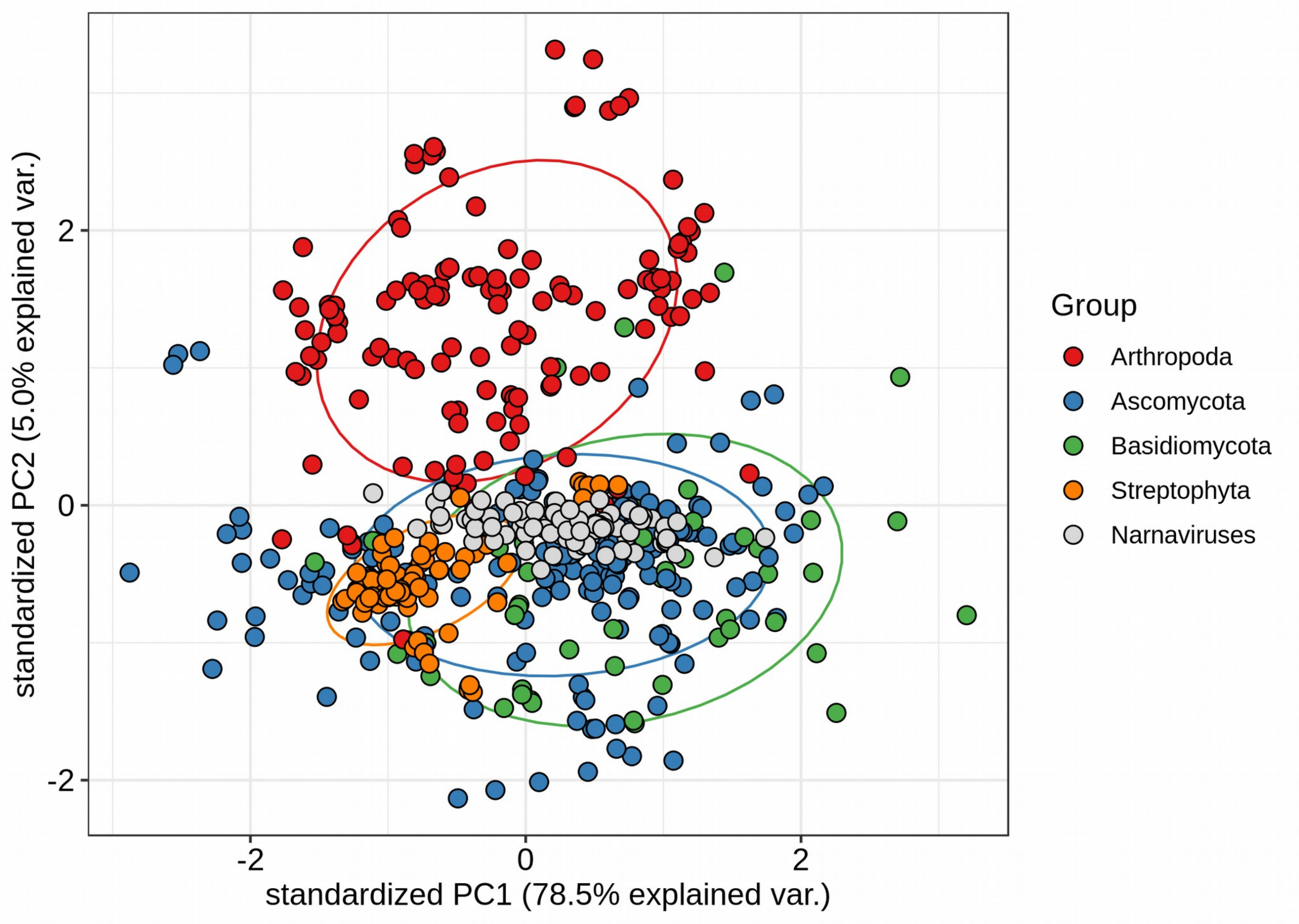


**Figure S5**. Principal component analysis of codon usage (per associated amino acid) in phyla that frequently co-occur with narnaviruses. Codon usage across NCBI RefSeq genomes was extracted from the latest release of the codon usage table database (CUTD). Ellipse contours are drawn for each group at a normal probability of 0.68.

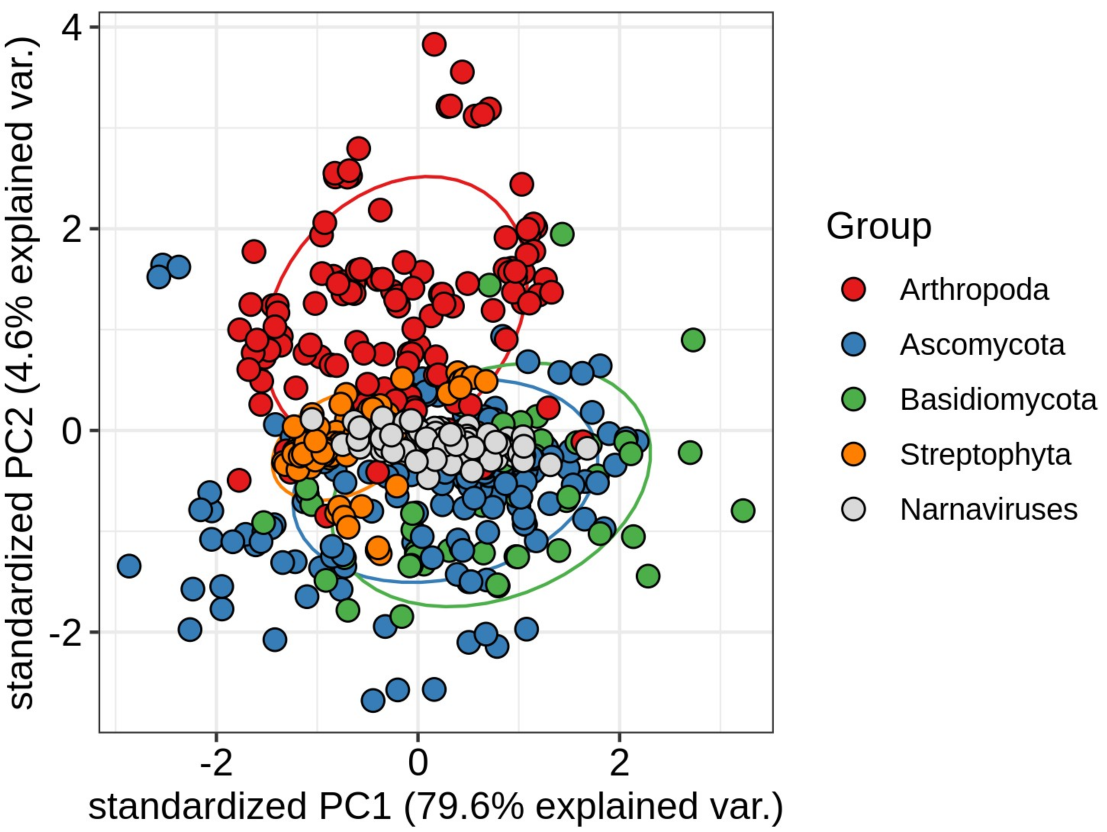


**Figure S6**. Principal component analysis of codon usage (per associated amino acid) in phyla that frequently co-occur with narnaviruses, excluding Leucine- and Serine-encoding codons. Codon usage across NCBI RefSeq genomes was extracted from the latest release of the codon usage table database (CUTD). Ellipse contours are drawn for each group at a normal probability of 0.68.
